# skater: An R package for SNP-based Kinship Analysis, Testing, and Evaluation

**DOI:** 10.1101/2021.07.21.453083

**Authors:** Stephen D. Turner, V. P. Nagraj, Matthew Scholz, Shakeel Jessa, Carlos Acevedo, Jianye Ge, August E. Woerner, Bruce Budowle

## Abstract

**Motivation:** SNP-based kinship analysis with genome-wide relationship estimation and IBD segment analysis methods produces results that often require further downstream processing and manipulation. A dedicated software package that consistently and intuitively implements this analysis functionality is needed.

**Results:** Here we present the skater R package for **S**NP-based **k**inship **a**nalysis, **t**esting, and **e**valuation with **R**. The skater package contains a suite of well-documented tools for importing, parsing, and analyzing pedigree data, performing relationship degree inference, benchmarking relationship degree classification, and summarizing IBD segment data.

**Availability:** The skater package is implemented as an R package and is released under the MIT license at https://github.com/signaturescience/skater. Documentation is available at https://signaturescience.github.io/skater.

## Introduction

Inferring familial relationships between individuals using genetic data is a common practice in population genetics, medical genetics, and forensics. There are multiple approaches to estimating relatedness between samples, including genome-wide measures, such as those implemented in Plink [1] or KING [2], and methods that rely on identity by descent (IBD) segment detection, such as GERMLINE [3], hap-IBD [4], and IBIS [5]. Recent efforts focusing on benchmarking these methods [6] have been aided by tools for simulating pedigrees and genome-wide SNP data [7]. Analyzing results from genome-wide SNP-based kinship analysis or comparing analyses to simulated data for benchmarking have to this point required writing one-off analysis functions or utility scripts that are seldom distributed with robust documentation, test suites, or narrative examples of usage. There is a need in the field for a well-documented software package with a consistent design and API that contains functions to assist with downstream manipulation, benchmarking, and analysis of SNP-based kinship assessment methods. Here we present the skater package for **S**NP-based **k**inship **a**nalysis, **t**esting, and **e**valuation with **R**.

## Methods

### Implementation

The skater package provides an intuitive collection of analysis and utility functions for SNP-based kinship analysis. Functions in the package include tools for importing, parsing, and analyzing pedigree data, performing relationship degree inference, benchmarking relationship degree classification, and summarizing IBD segment data, described in full in the *Use Cases* section below. The package adheres to “tidy” data analysis principles, and builds upon the tools released under the tidyverse R ecosystem [8].

The skater package is hosted in the Comprehensive R Archive Network (CRAN) which is the main repository for R packages: http://CRAN.R-project.org/package=skater. Users can install skater in R by executing the following code:

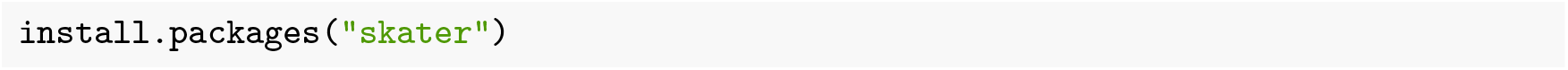

Alternatively, the development version of skater is available on GitHub at https://github.com/signaturescience/skater. The development version may contain new features which are not yet available in the version hosted on CRAN. This version can be installed using the `install_github()` function in the devtools package:

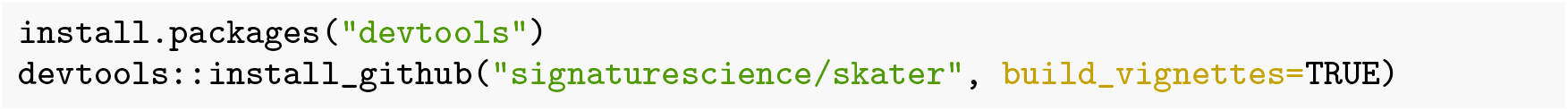

When installing skater, other packages which skater depends on are automatically installed, including magritr, tibble, dplyr, tidyr, readr, purrr, kinship2, corrr, rlang, and others.

### Operation

Minimal system requirements for installing and using skater include R (version 3.0.0 or higher) and several tidyverse packages [8] that many R users will already have installed. Use cases are demonstrated in detail below. In summary, the skater package has functions for:

- Reading in various output files produced by commonly used tools in SNP-based kinship analysis
- Pedigree parsing, manpulation, and analysis
- Relationship degree inference
- Benchmarking and assessing relationship classification accuracy
- IBD segment analysis post-processing

A comprehensive reference for all the functions in the skater package is available at https://signaturescience.github.io/skater/.

#### Use Cases

The `skater` package provides a collection of analysis and utility functions for **S**NP-based **k**inship **a**nalysis, **t**esting, and **e**valuation as an **R** package. Functions in the package include tools for working with pedigree data, performing relationship degree inference, assessing classification accuracy, and summarizing IBD segment data.

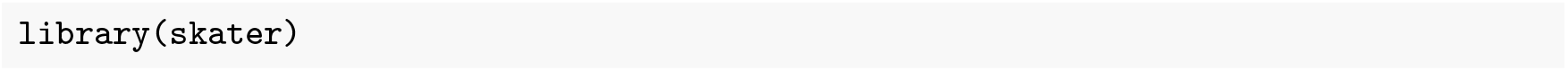

### Pedigree parsing, manipulation, and analysis

Pedigrees define familial relationships in a hierarchical structure. One of the common formats used by PLINK [1] and other genetic analysis tools is the `.fam` file. A `.fam` file is a tabular format with one row per individual and columns for unique IDs of the mother, father, and the family unit. The package includes `read_fam()` to read files in this format:

Family structures imported from `.fam` formated files can then be translated to the `pedigree` structure used by the `kinship2` package [9]. The “fam” format may include multiple families, and the `fam2ped()` function will collapse them all into a `tibble` with one row per family:

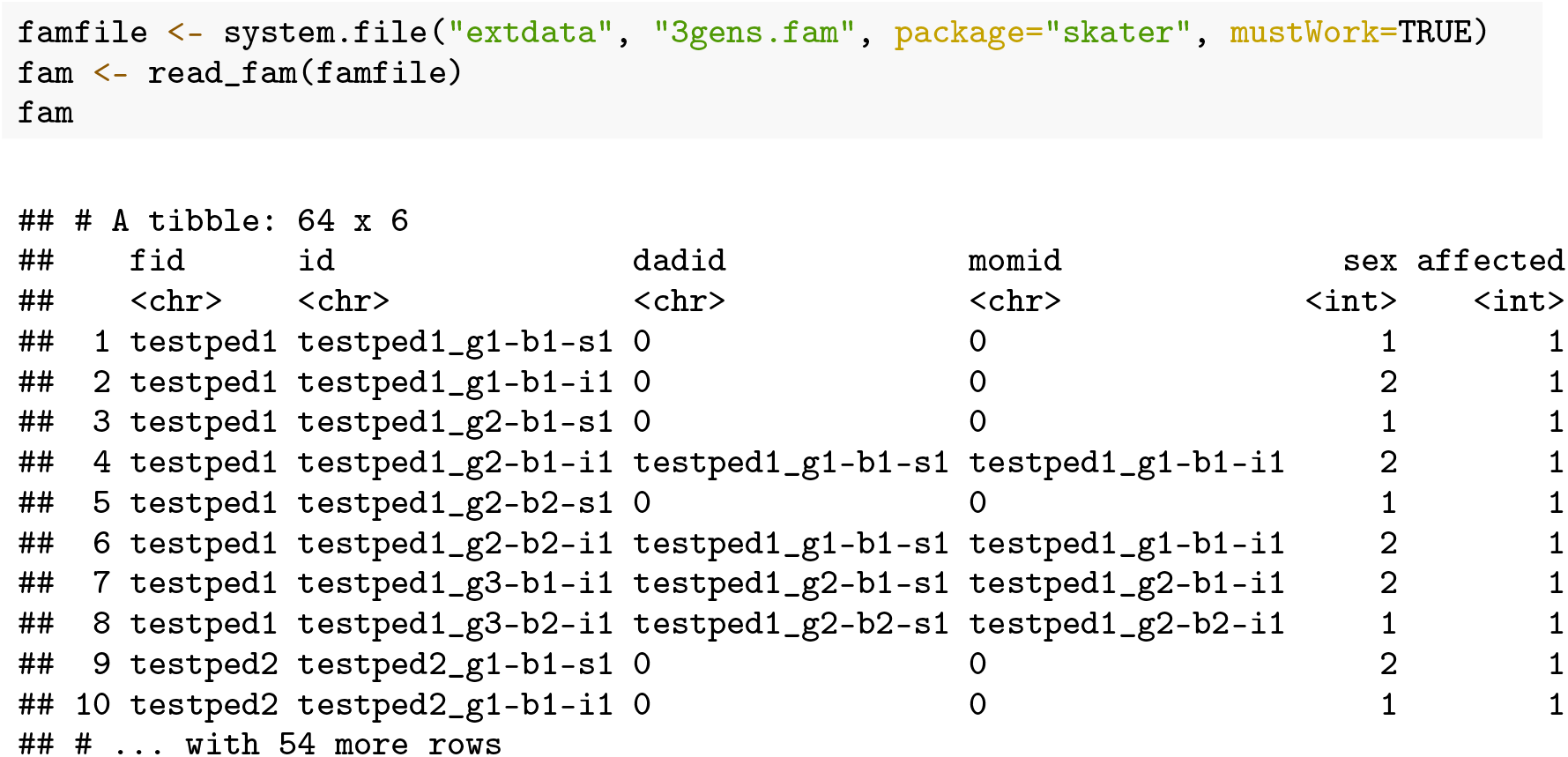

In the example above, the resulting `tibble` is nested by family ID. The `data` column contains the individual family information, while the `ped` column contains the pedigree object for that family. Using standard tidyverse operations, the resulting tibble can be unnested for any particular family:

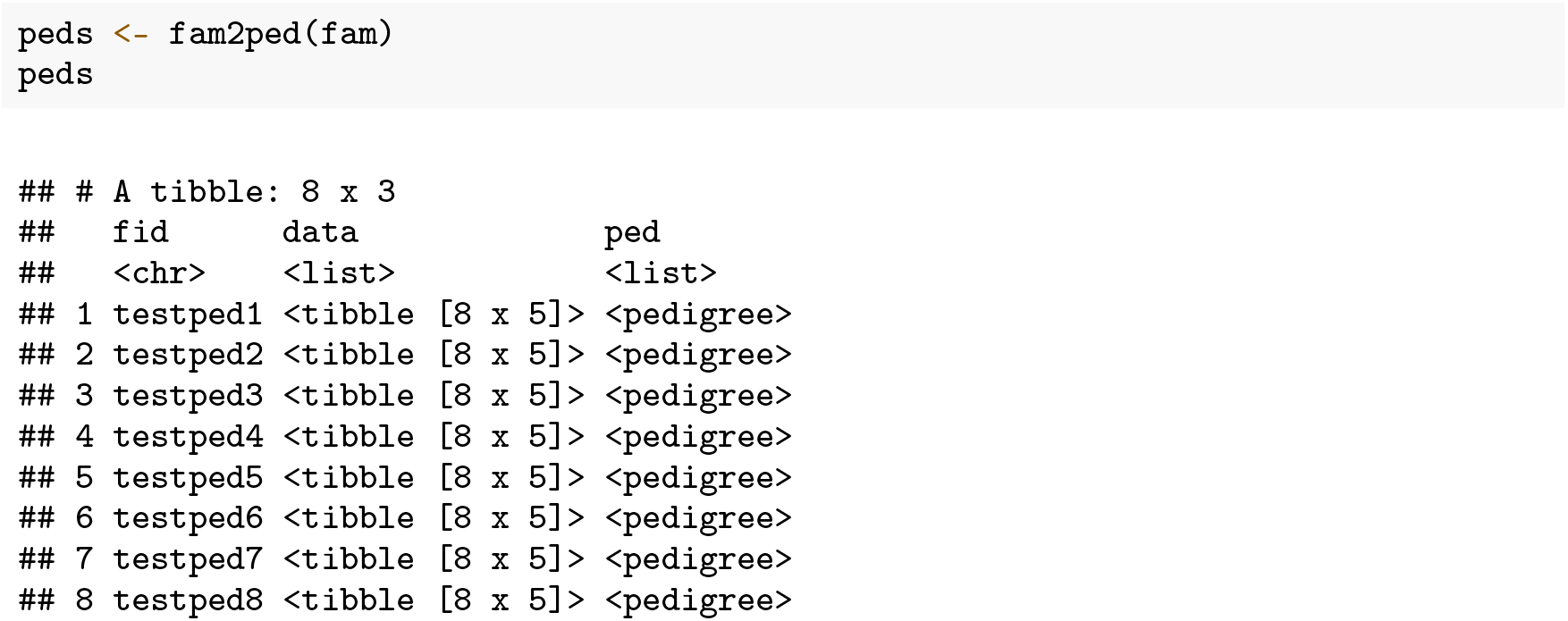

A single pedigree can also be inspected or visualized (standard base R plot arguments such as `mar` or `cex` can be used to adjust aesthetics):

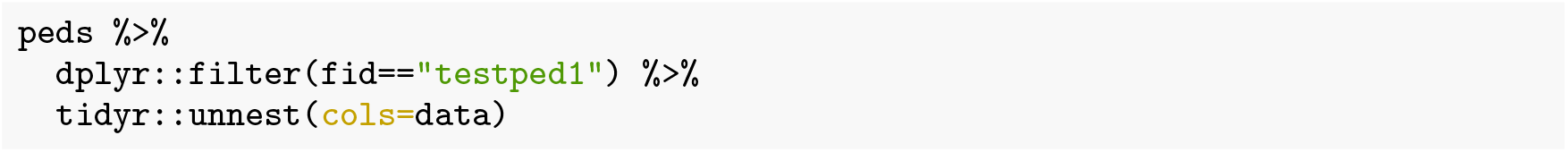
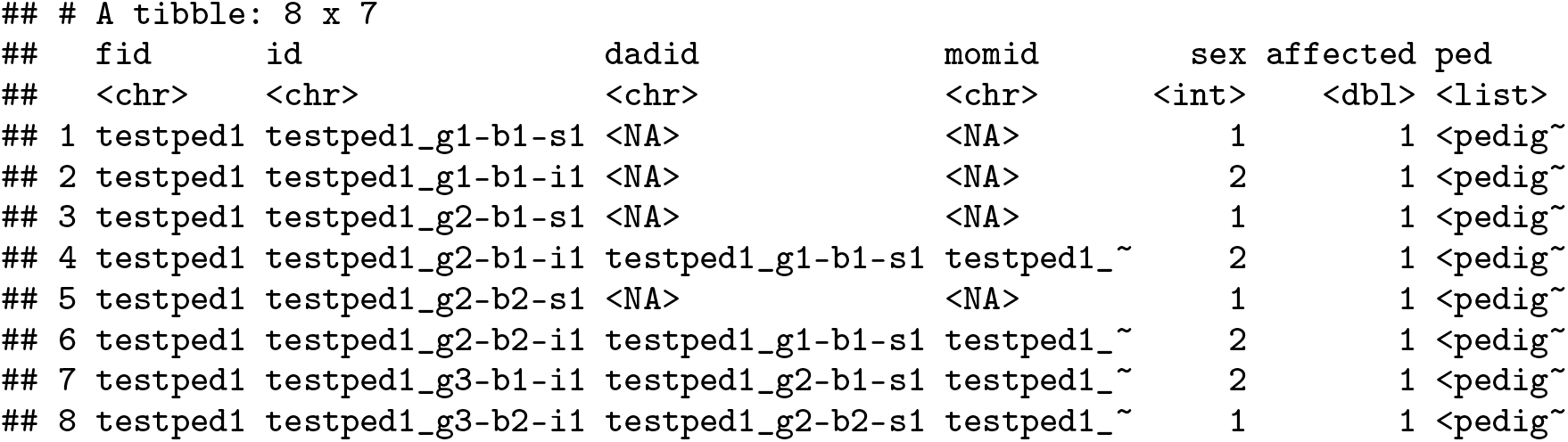

The `plot_pedigree()` function from `skater` will iterate over a list of pedigree objects, writing a multi-page PDF, with each page containing a pedigree from family:

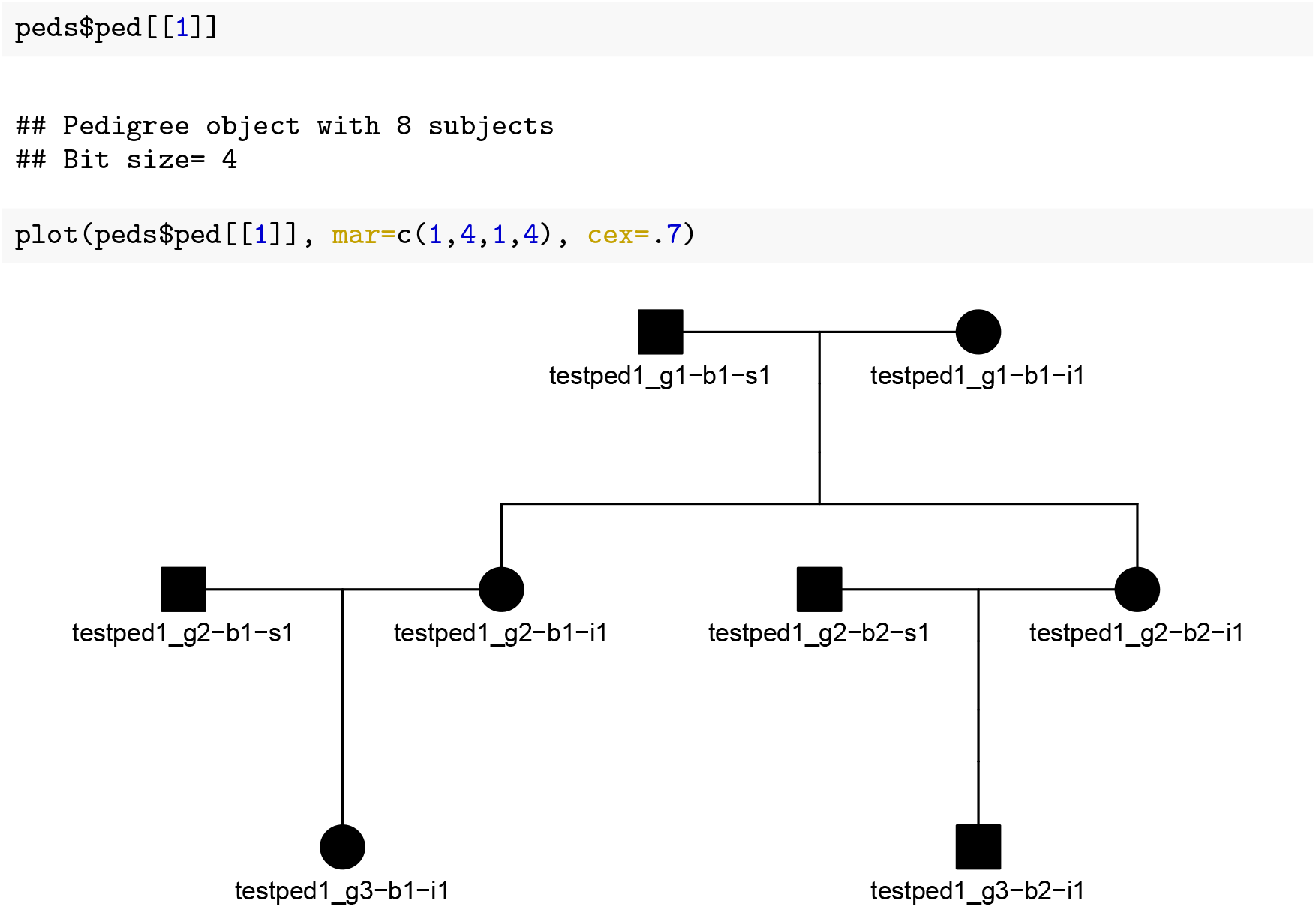

The `ped2kinpair()` function takes a pedigree object and produces a pairwise list of relationships between all individuals in the data with the expected kinship coefficients for each pair.

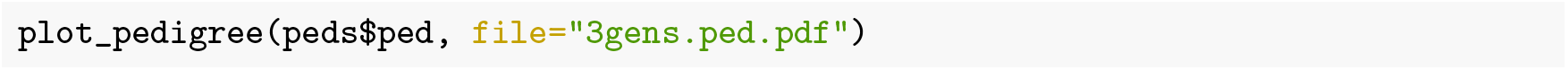

The function can be run on a single family:

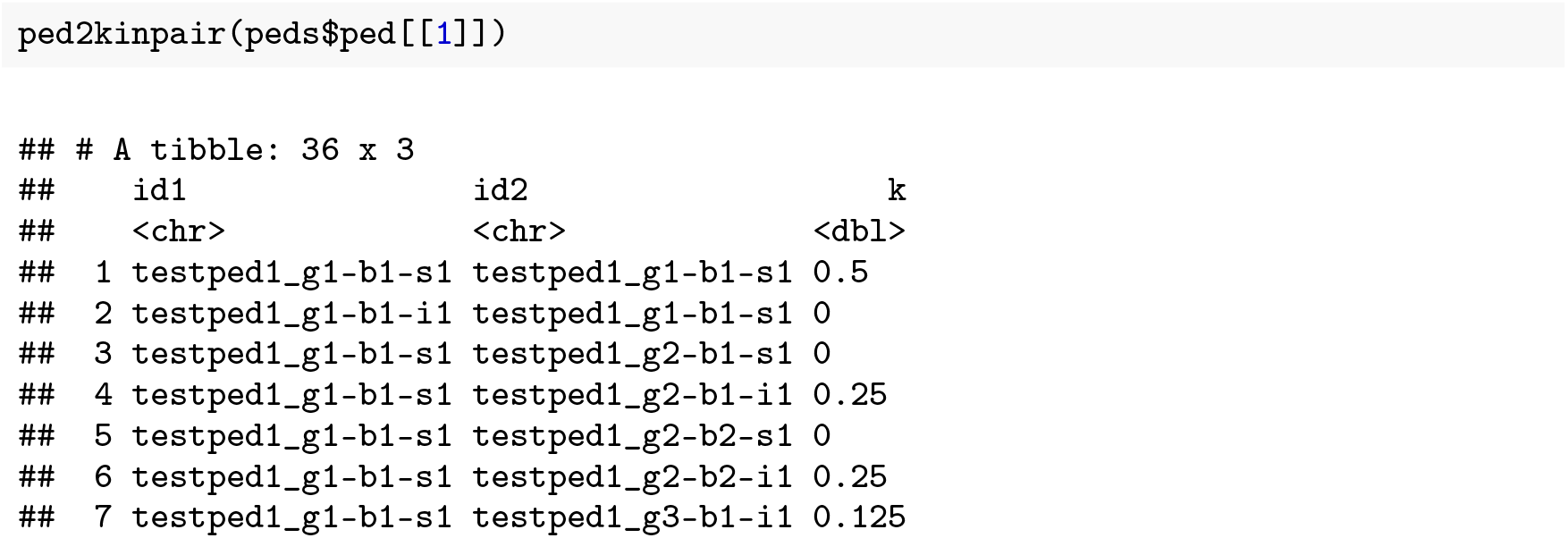
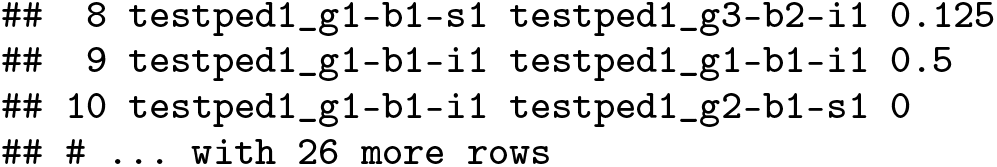

This function can also be mapped over all families in the pedigree:

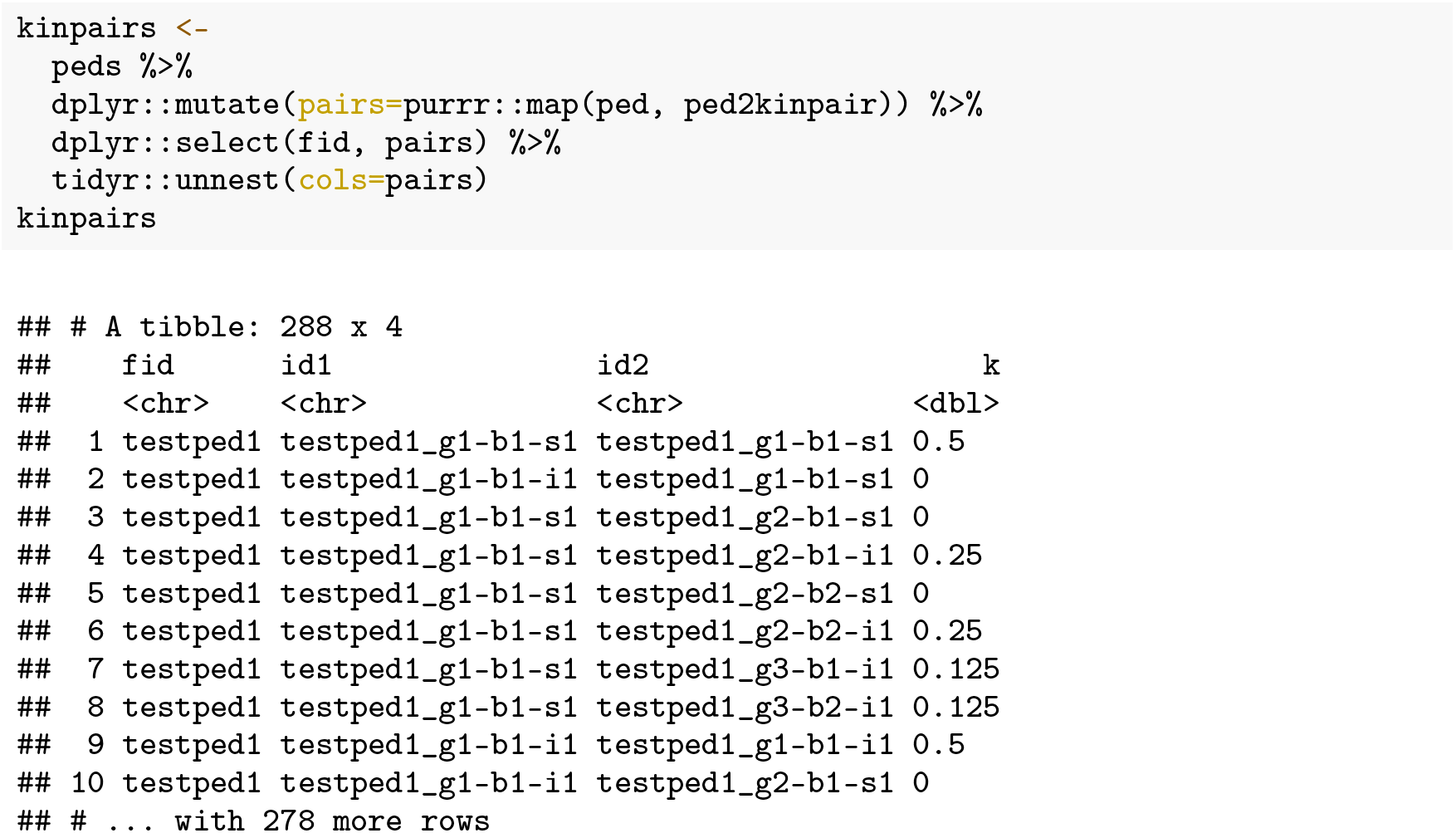

Note that this maps `ped2kinpair()` over all `ped` objects in the input `tibble`, and that relationships are not shown for between-family relationships.

### Relationship degree inference and benchmarking

The skater package includes functions to translate kinship coefficients to relationship degrees. The kinship coefficients could come from `ped2kinpair()` or other kinship estimation software.

The `dibble()` function creates a **d**egree **i**nference `tibble`, with degrees up to the specified `max_degree` (default=3), expected kinship coefficient, and lower (`l`) and upper (`u`) inference ranges as defined in Manichaikul et al. [2]. Degree 0 corresponds to self / identity / monozygotic twins, with an expected kinship coefficient of 0.5, with inference range >=0.354. Anything beyond the maximum degree resolution is considered unrelated (degree `NA`). Note also that while the theoretical upper boundary for the kinship coefficient is 0.5, the inference range for 0-degree (same person or identical twins) extends to 1 to allow for floating point arithmetic and stochastic effects resulting in kinship coefficients above 0.5.

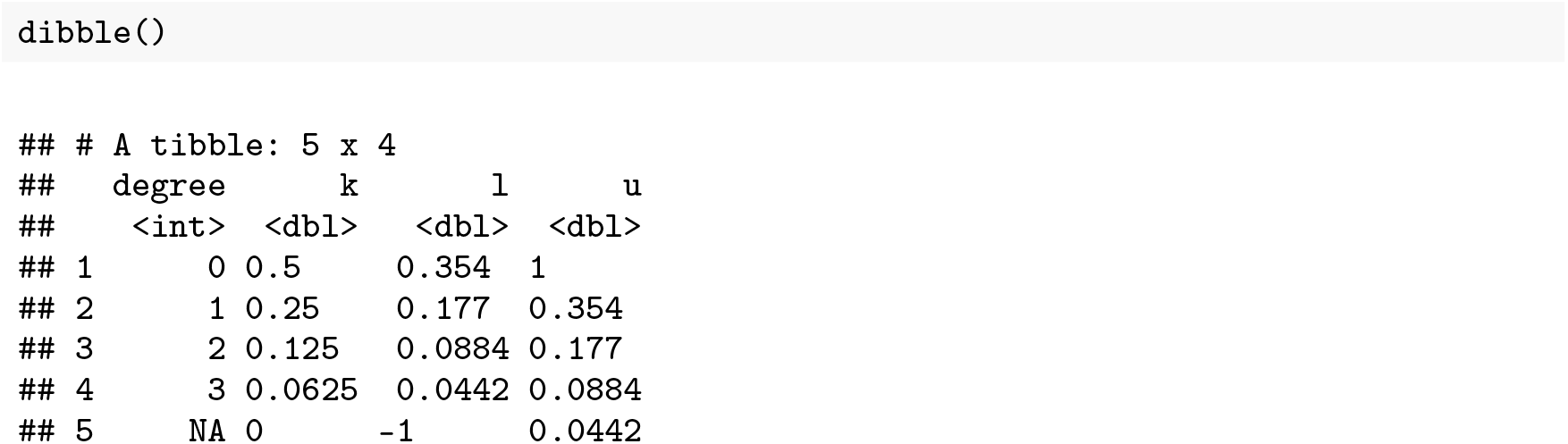

The degree inference `max_degree` default is 3. Change this argument to allow more granular degree inference ranges:

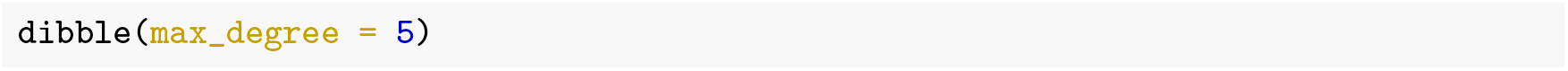
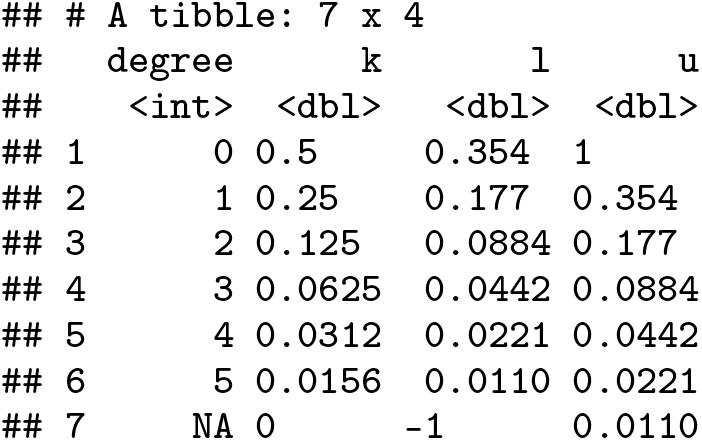

Note that the distance between relationship degrees becomes smaller as the relationship degree becomes more distant. The `dibble()` function will emit a warning with `max_degree` >=10, and will stop with an error at >=12.

The `kin2degree()` function infers the relationship degree given a kinship coefficient and a `max_degree` up to which anything more distant is treated as unrelated. Example first degree relative:

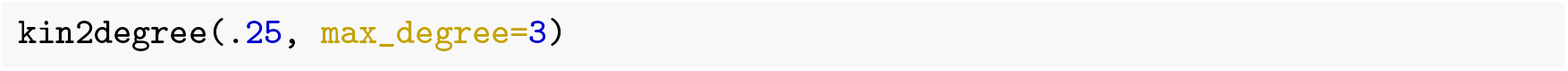

\## [1] 1

Example 4th degree relative, but using the default max_degree resolution of 3:

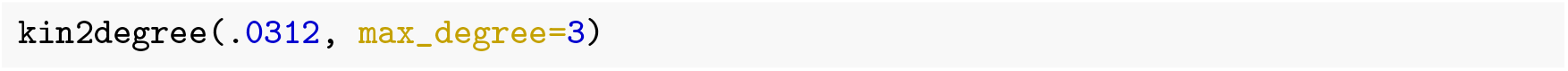

\## [1] 1

Example 4th degree relative, but increasing the degree resolution:

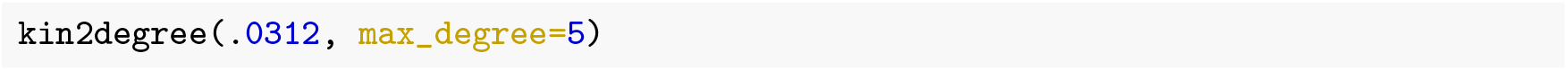

\## [1] 1

The `kin2degree()` function is vectorized over values of `k`, so it can be used inside of a `mutate` on a `tibble` of kinship coefficients:

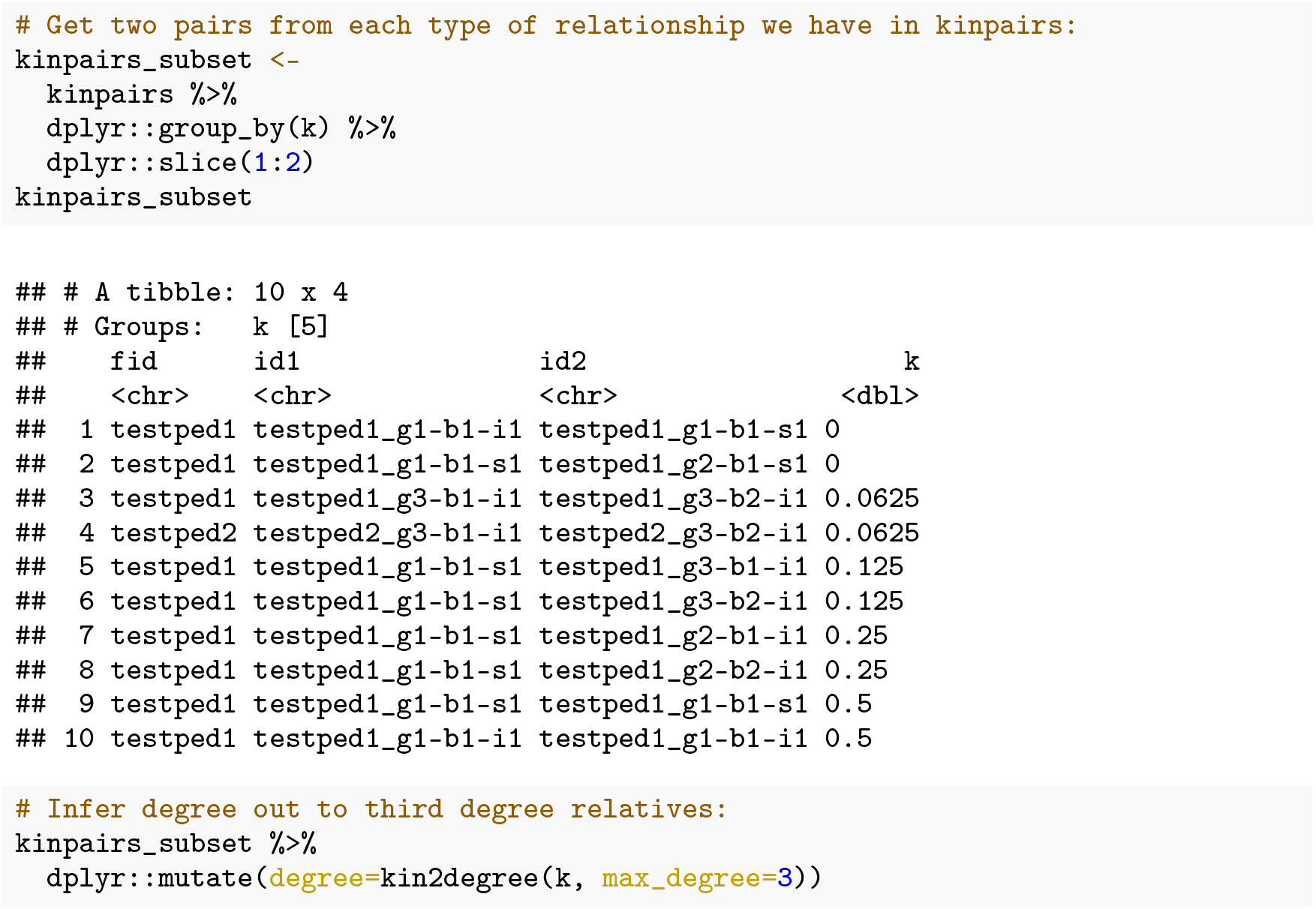
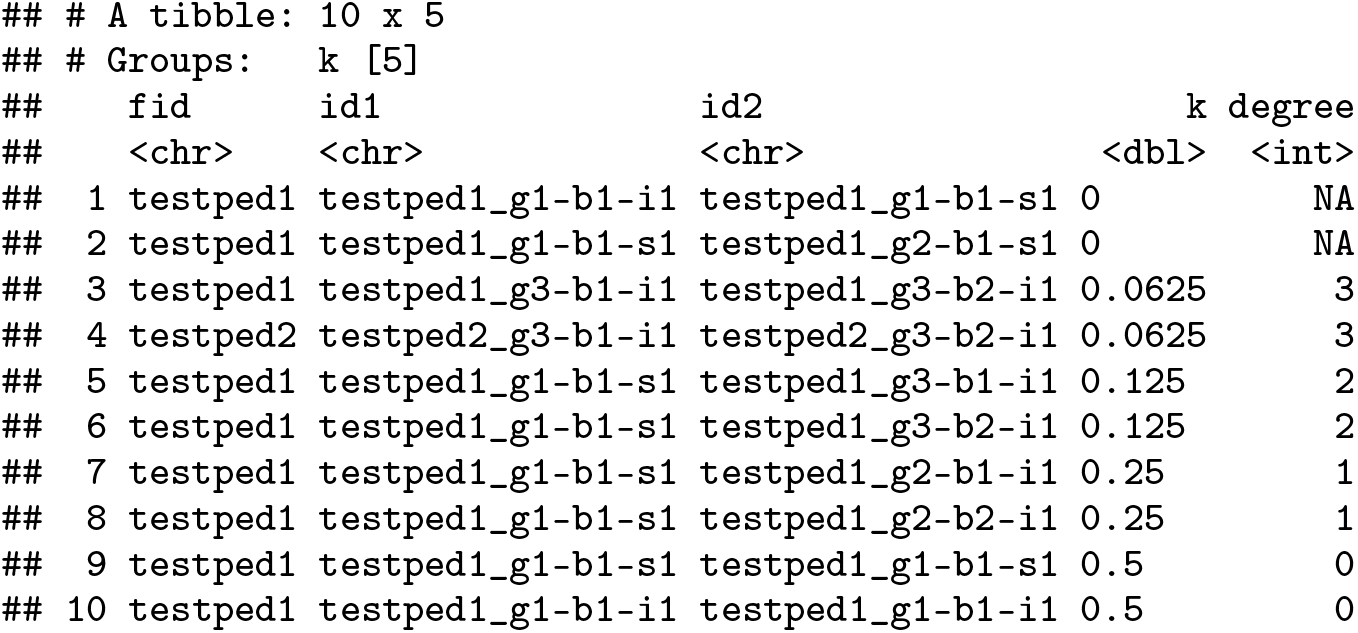

### Benchmarking Degree Classification

Once estimated kinship is converted to degree, it may be of interest to compare the inferred degree to truth. When aggregated over many relationships and inferences, this approach can help benchmark performance of a particular kinship analysis method.

The skater package adapts a `confusion_matrix()` function from Clark [10] to provide standard contingency table metrics (e.g. sensitivity, specificity, PPV, precision, recall, F1, etc.) with a new reciprocal RMSE (R-RMSE) metric. The `confusion_matrix()` function on its own outputs a list with four objects:

1. A `tibble` with calculated accuracy, lower and upper bounds, the guessing rate and p-value of the accuracy vs. the guessing rate.
2. A `tibble` with contingency table statistics calculated for each class. Details on the statistics calculated for each class can be reviewed on the help page for `?confusion_matrix`.
3. A `matrix` with the contingency table object itself.
4. A `vector` with the reciprocal RMSE (R-RMSE). The R-RMSE represents an alternative to classification accuracy when benchmarking relationship degree estimation and is calculated using the formula in (1). Taking the reciprocal of the target and predicted degree results in larger penalties for more egregious misclassifications (e.g., classifying a first-degree relative pair as second degree) than misclassifications at more distant relationships (e.g., misclassifying a fourth-degree relative pair as fifth-degree). The +0.5 adjustment prevents division-by-zero when a 0th-degree (identical) relative pair is introduced.

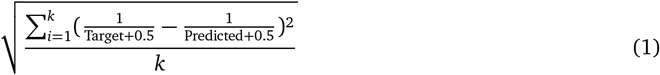

To illustrate the usage, this example will start with the `kinpairs` data from above and randomly flip ∼20% of the true relationship degrees:

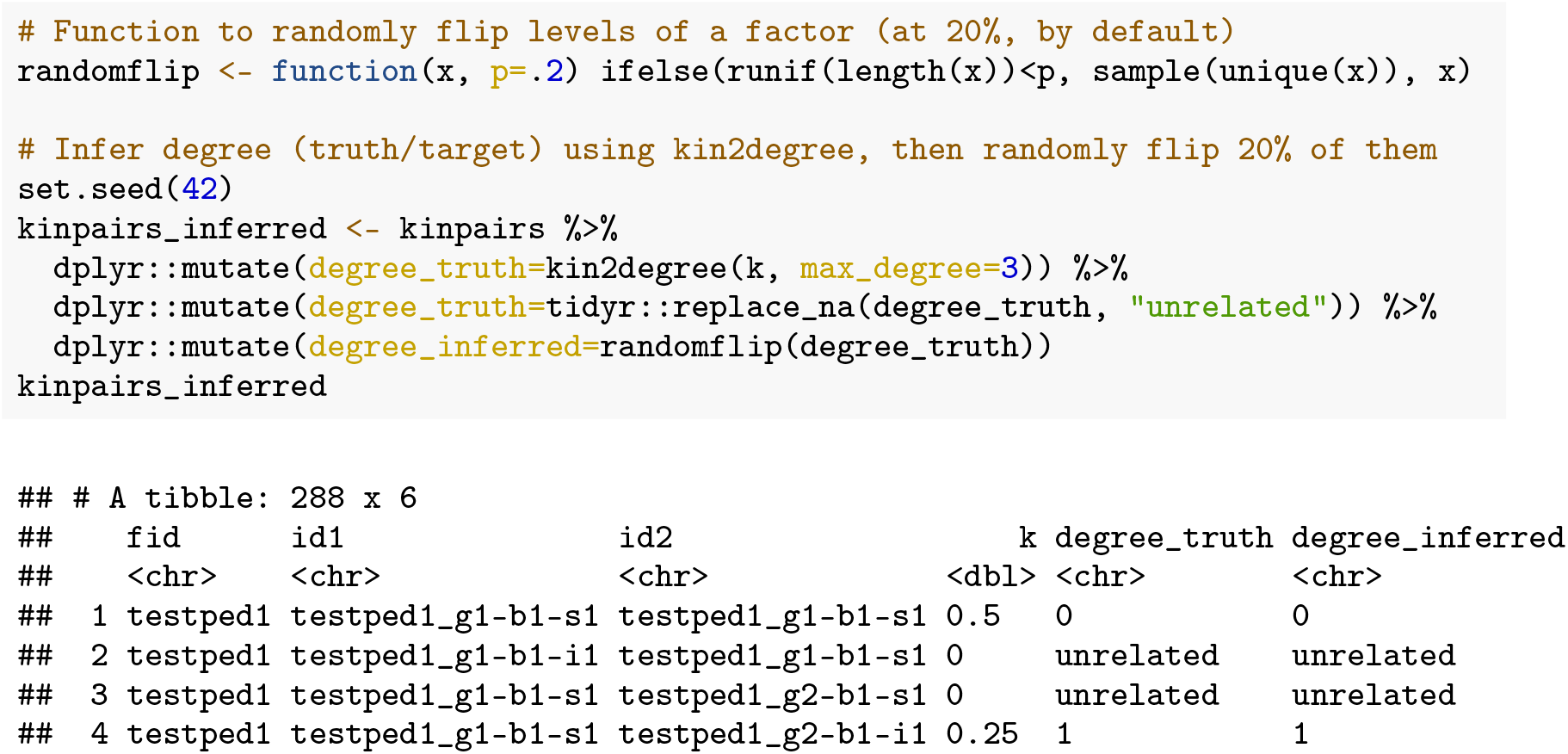
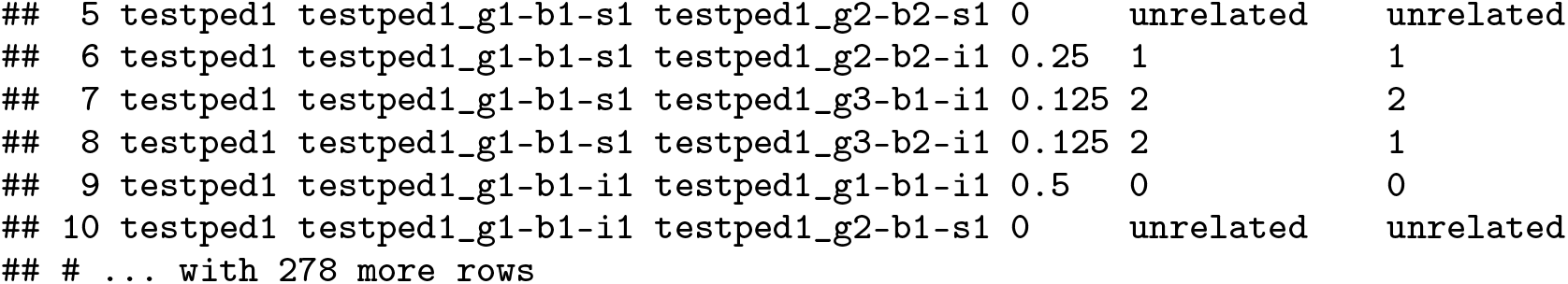

Next, running the `confusion_matrix()` function will return all four objects noted above:

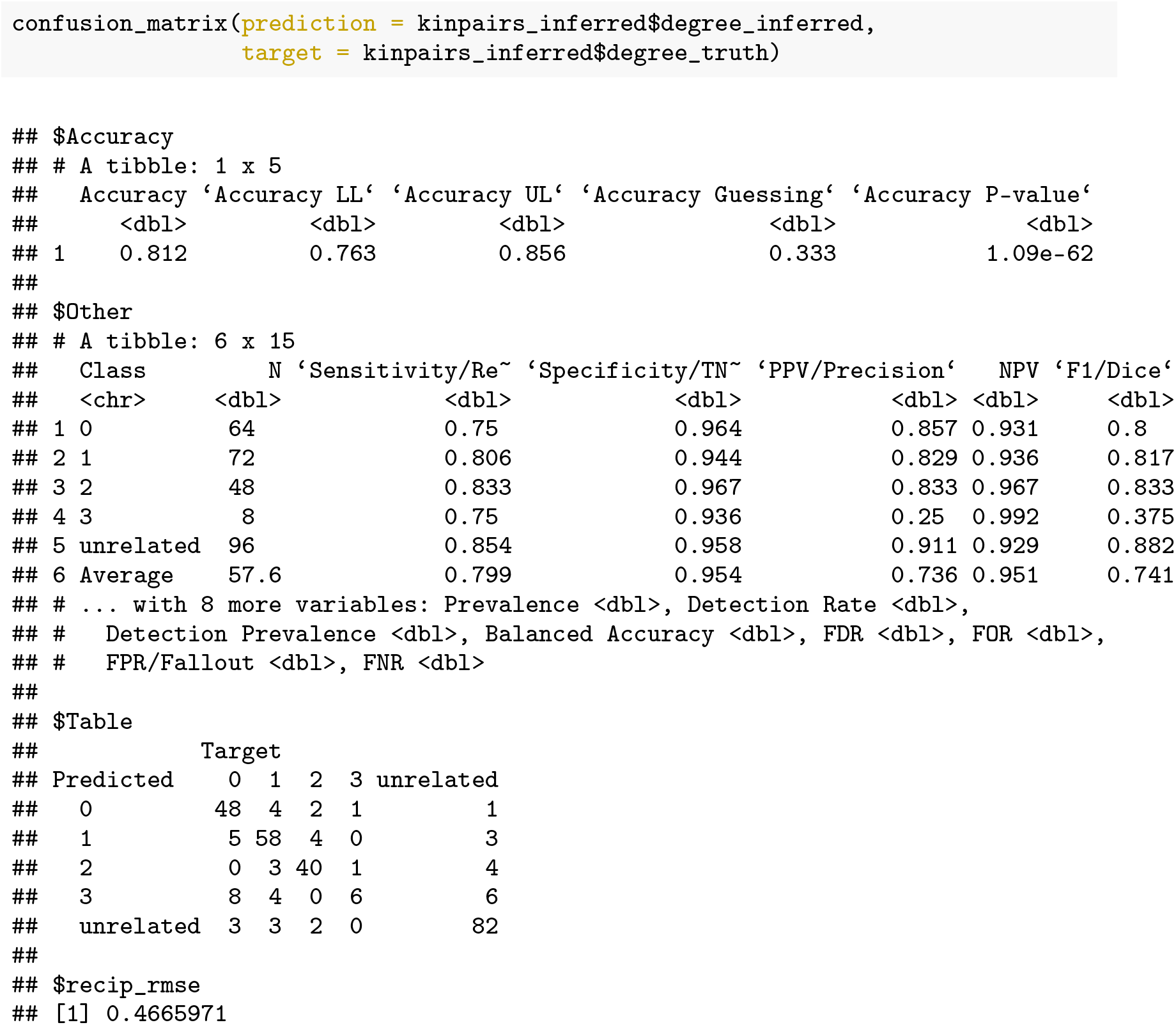

Standard tidyverse functions such as `purrr::pluck()` can be used to isolate just the contingency table:

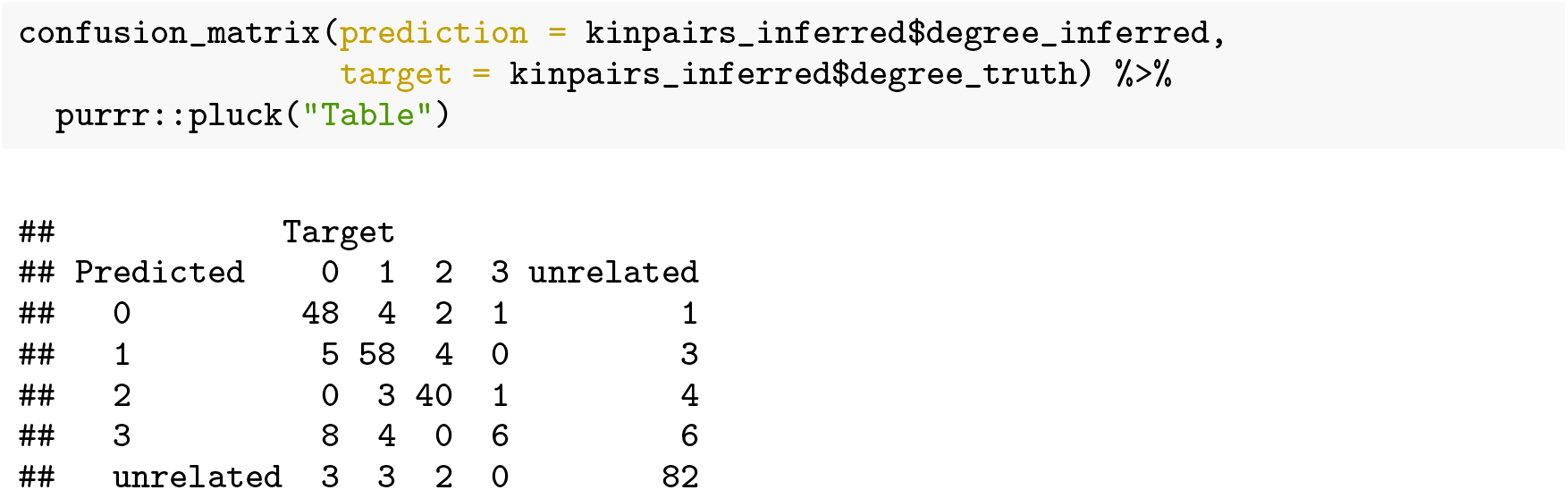

The `confusion_matrix()` function includes an argument to output in a tidy (`longer=TRUE`) format, and the example below illustrates how to spread contingency table statistics by class:

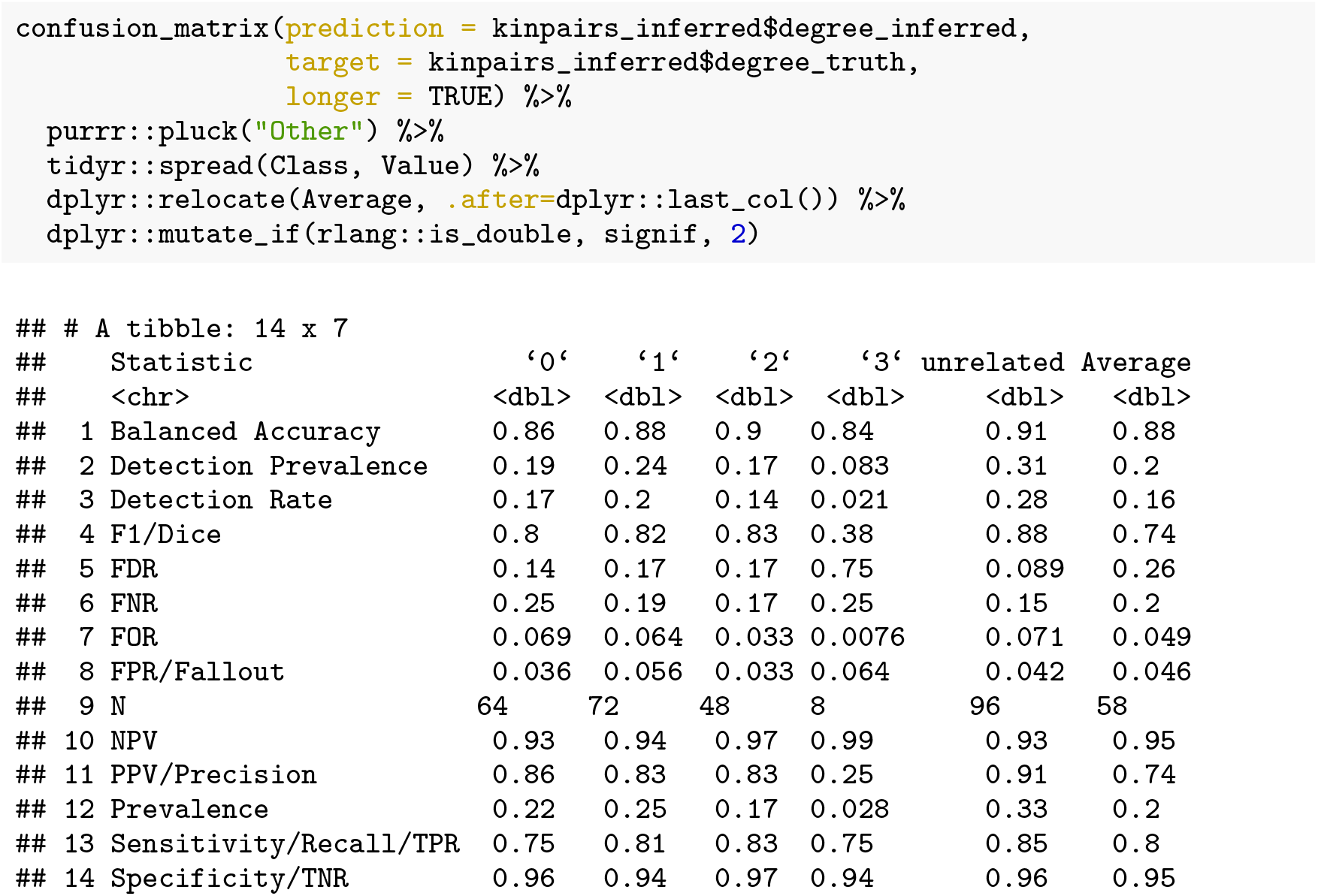

### IBD segment analysis

Tools such as hap-IBD [4], and IBIS [5] detect shared IBD segments between individuals. The skater package includes functionality to take those IBD segments, compute shared genomic centimorgan (cM) length, and converts that shared cM to a kinship coefficient. In addition to inferred segments, these functions can estimate “truth” kinship from simulated IBD segments [7]. The `read_ibd()` function reads pairwise IBD segments from IBD inference tools and from simulated IBD segments. The `read_map()` function reads in genetic map in a standard format which is required to translate the total centimorgans shared IBD to a kinship coefficient using the `ibd2kin()` function. See `?read_ibd` and `?read_map` for additional details on expected format.

The `read_ibd()` function reads in the pairwise IBD segment format. Input to this function can either be inferred IBD segments from hap-IBD (`source=“hapibd”`) or simulated segments (`source=“pedsim”`). The first example below uses data in the `hap-ibd` output format:

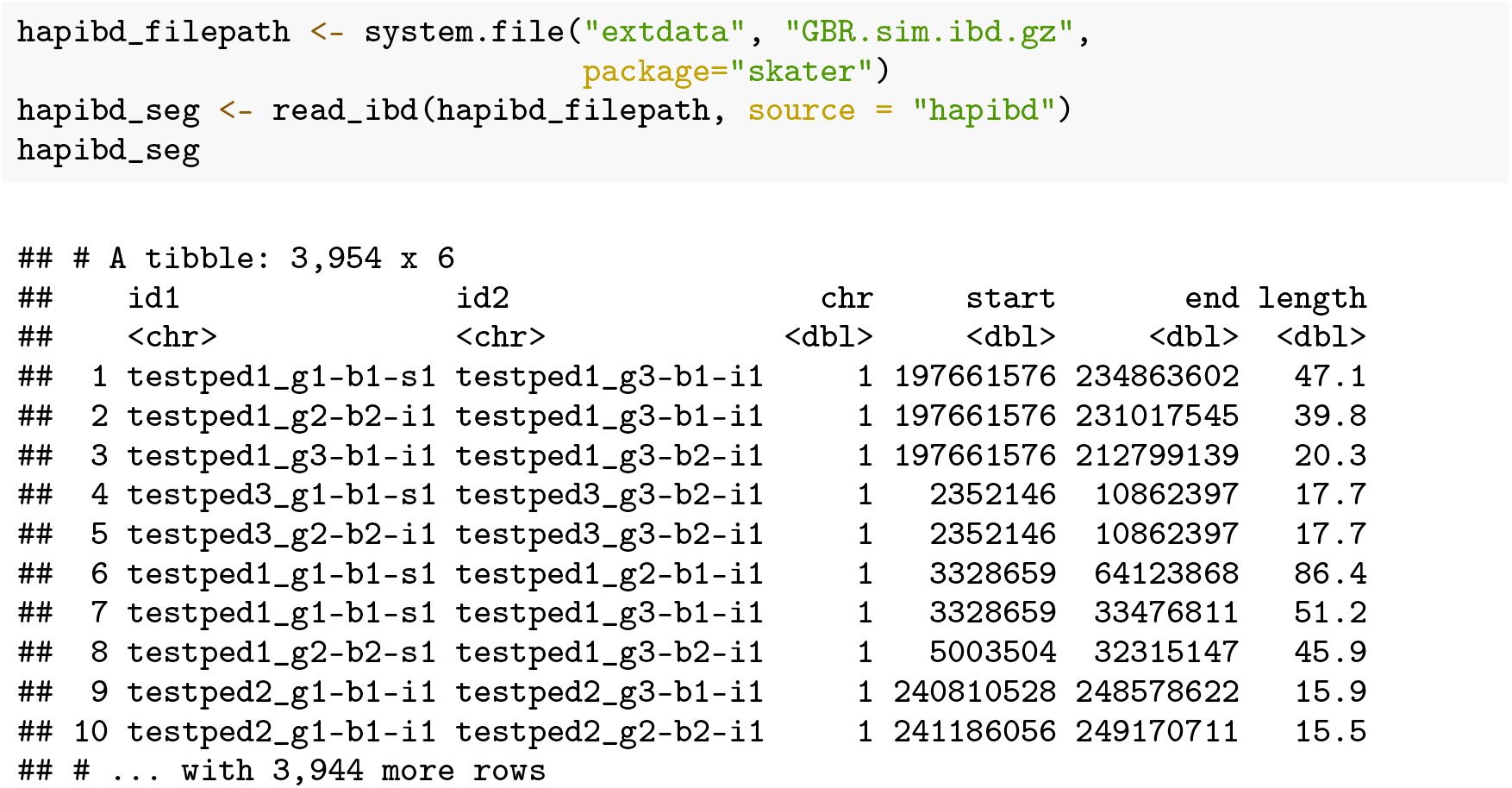

In order to translate the shared genomic cM length to a kinship coefficient, a genetic map must first be read in with `read_map()`. Software for IBD segment inference and simulation requires a genetic map. The map loaded for kinship estimation should be the same one used for creating the shared IBD segment output. The example below uses a minimal genetic map that ships with `skater`:

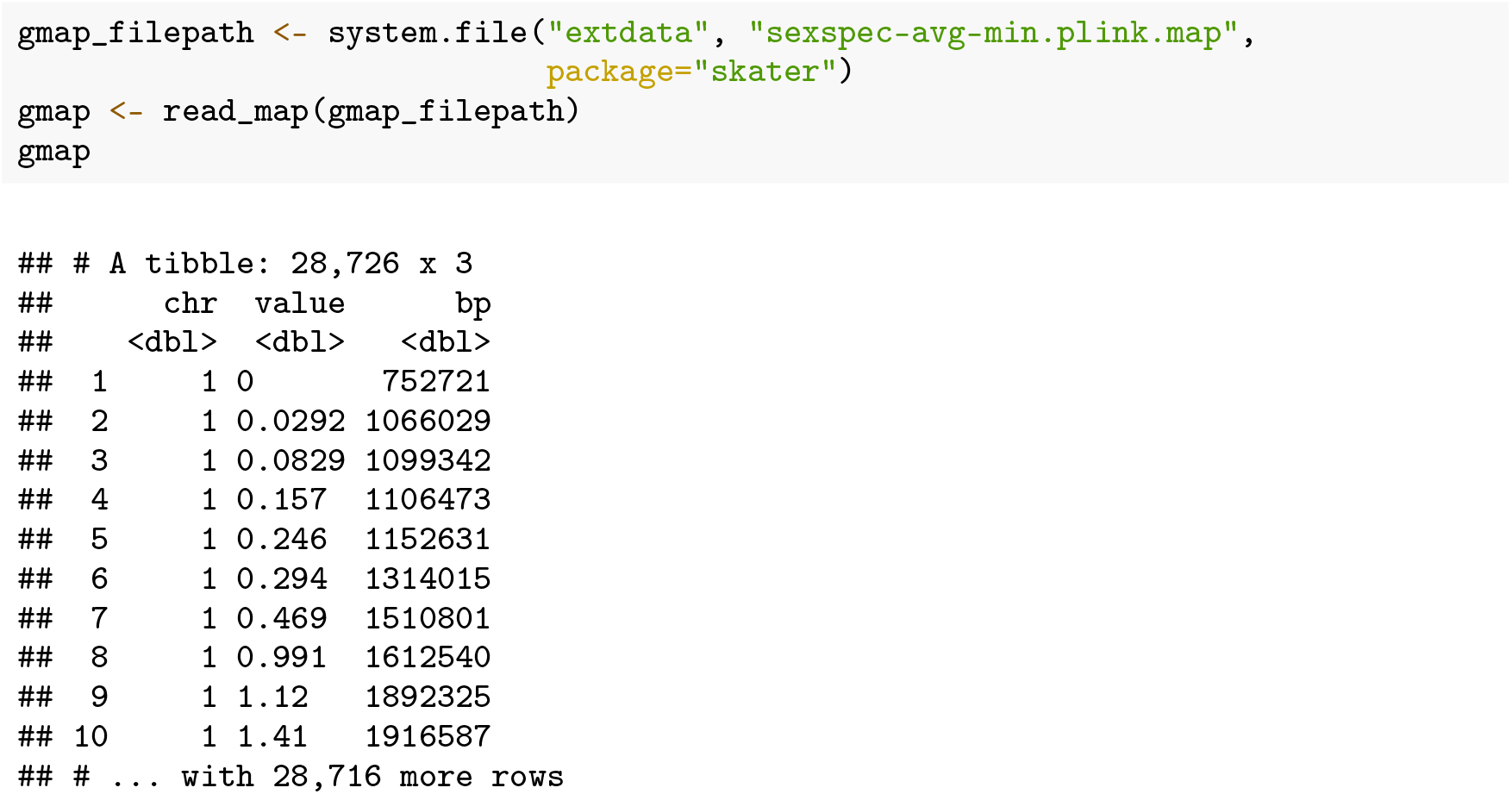

The `ibd2kin()` function takes the segments and map file and outputs a `tibble` with one row per pair of individuals and columns for individual 1 ID, individual 2 ID, and the kinship coefficient for the pair:

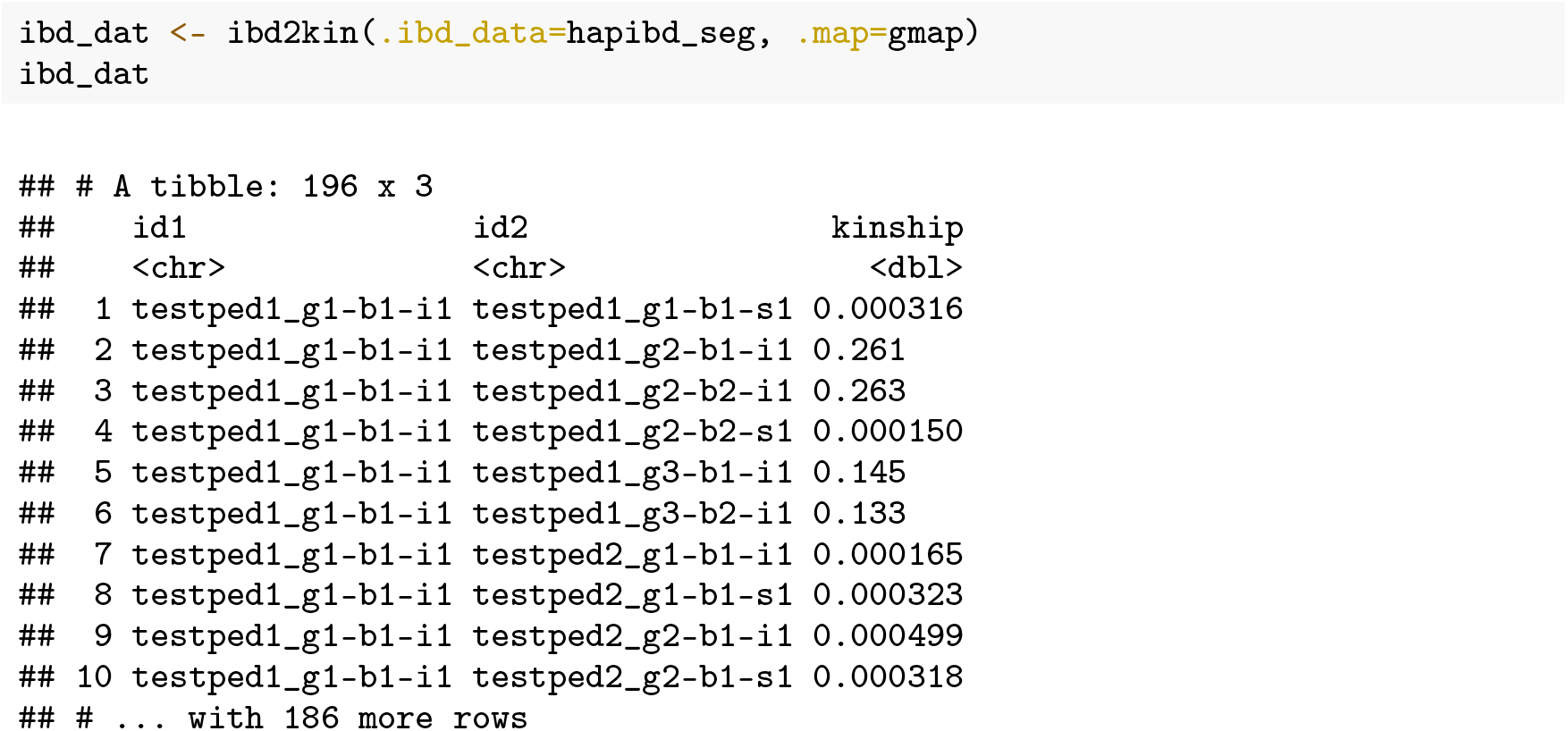

## Summary

The skater R package provides a robust software package for data import, manipulation, and analysis tasks typically encountered when working with SNP-based kinship analysis tools. All package functions are internally documented with examples, and the package contains a vignette demonstrating usage, inputs, outputs, and interpretation of all key functions. The package contains internal tests that are automatically run with continuous integration via GitHub Actions whenever the package code is updated. The skater package is permissively licensed (MIT) and is easily extensible to accommodate outputs from new genome-wide relatedness and IBD segment methods as they become available.

## Software availability

1. Software available from: http://CRAN.R-project.org/package=skater.
2. Source code available from: https://github.com/signaturescience/skater.
3. Archived source code at time of publication: https://doi.org/10.5281/zenodo.5761996.
4. Software license: MIT License.

## Author information

SDT, VPN, and MBS developed the R package.

All authors contributed to method development.

SDT wrote the first draft of the manuscript.

All authors assisted with manuscript revision.

All authors read and approved the final manuscript.

## References

[1] Shaun Purcell, Benjamin Neale, Kathe Todd-Brown, Lori Thomas, Manuel A. R. Ferreira, David Bender, Julian Maller, Pamela Sklar, Paul I. W. de Bakker, Mark J. Daly, and Pak C. Sham. PLINK: A Tool Set for Whole-Genome Association and Population-Based Linkage Analyses. The American Journal of Human Genetics, 81(3):559–575, September 2007. ISSN 0002-9297. doi: 10.1086/519795.

[2] Ani Manichaikul, Josyf C. Mychaleckyj, Stephen S. Rich, Kathy Daly, Michèle Sale, and Wei-Min Chen. Robust relationship inference in genome-wide association studies. Bioinformatics (Oxford, England), 26(22):2867–2873, November 2010. ISSN 1367-4811. doi: 10.1093/bioinformatics/btq559.

[3] Alexander Gusev, Jennifer K. Lowe, Markus Stoffel, Mark J. Daly, David Altshuler, Jan L. Breslow, Jeffrey M. Friedman, and Itsik Pe’er. Whole population, genome-wide mapping of hidden relatedness. Genome Research, 19(2):318–326, February 2009. ISSN 1088-9051, 1549-5469. doi: 10.1101/gr.081398.108.

[4] Ying Zhou, Sharon R. Browning, and Brian L. Browning. A Fast and Simple Method for Detecting Identity-by-Descent Segments in Large-Scale Data. The American Journal of Human Genetics, 106(4):426–437, April 2020. ISSN 0002-9297. doi: 10.1016/j.ajhg.2020.02.010.

[5] Daniel N. Seidman, Sushila A. Shenoy, Minsoo Kim, Ramya Babu, Ian G. Woods, Thomas D. Dyer, Donna M. Lehman, Joanne E. Curran, Ravindranath Duggirala, John Blangero, and Amy L. Williams. Rapid, Phase-free Detection of Long Identity-by-Descent Segments Enables Effective Relationship Classification. American Journal of Human Genetics, 106 (4):453–466, April 2020. ISSN 1537-6605. doi: 10.1016/j.ajhg.2020.02.012.

[6] Monica D. Ramstetter, Thomas D. Dyer, Donna M. Lehman, Joanne E. Curran, Ravindranath Duggirala, John Blangero, Jason G. Mezey, and Amy L. Williams. Benchmarking Relatedness Inference Methods with Genome-Wide Data from Thousands of Relatives. Genetics, 207(1):75–82, September 2017. ISSN 0016-6731, 1943-2631. doi: 10.1534/genetics.117.1122.

[7] Madison Caballero, Daniel N. Seidman, Ying Qiao, Jens Sannerud, Thomas D. Dyer, Donna M. Lehman, Joanne E. Curran, Ravindranath Duggirala, John Blangero, Shai Carmi, and Amy L. Williams. Crossover interference and sex-specific genetic maps shape identical by descent sharing in close relatives. PLOS Genetics, 15(12):e1007979, December 2019. ISSN 1553-7404. doi: 10.1371/journal.pgen.1007979.

[8] Hadley Wickham, Mara Averick, Jennifer Bryan, Winston Chang, Lucy D’Agostino McGowan, Romain François, Garrett Grolemund, Alex Hayes, Lionel Henry, Jim Hester, Max Kuhn, Thomas Lin Pedersen, Evan Miller, Stephan Milton Bache, Kirill Müller, Jeroen Ooms, David Robinson, Dana Paige Seidel, Vitalie Spinu, Kohske Takahashi, Davis Vaughan, Claus Wilke, Kara Woo, and Hiroaki Yutani. Welcome to the tidyverse. Journal of Open Source Software, 4(43):1686, 2019. doi: 10.21105/joss.01686. URL https://doi.org/10.21105/joss.01686.

[9] Jason P. Sinnwell, Terry M. Therneau, and Daniel J. Schaid. The kinship2 R Package for Pedigree Data. Human heredity, 78(2):91–93, 2014. ISSN 0001-5652. doi: 10.1159/000363105.

[10] Michael Clark. https://github.com/m-clark/confusionmatrix, January 2021.

